# A Rigorous Multi-Laboratory Study of Known PDAC Biomarkers Identifies Increased Sensitivity and Specificity Over CA19-9 Alone

**DOI:** 10.1101/2024.05.22.595399

**Authors:** Brian Haab, Lu Qian, Ben Staal, Maneesh Jain, Johannes Fahrmann, Christine Worthington, Denise Prosser, Liudmila Velokokhatnaya, Camden Lopez, Runlong Tang, Mark W. Hurd, Gopalakrishnan Natarajan, Sushil Kumar, Lynette Smith, Sam Hanash, Surinder K. Batra, Anirban Maitra, Anna Lokshin, Ying Huang, Randall E. Brand

## Abstract

A blood test that enables surveillance for early-stage pancreatic ductal adenocarcinoma (PDAC) is an urgent need. Independent laboratories have reported PDAC biomarkers that could improve biomarker performance over CA19-9 alone, but the performance of the previously reported biomarkers in combination is not known. Therefore, we conducted a coordinated case/control study across multiple laboratories using common sets of blinded training and validation samples (132 and 295 plasma samples, respectively) from PDAC patients and non-PDAC control subjects representing conditions under which surveillance occurs. We analyzed the training set to identify candidate biomarker combination panels using biomarkers across laboratories, and we applied the fixed panels to the validation set. The panels identified in the training set, CA19-9 with CA199.STRA, LRG1, TIMP-1, TGM2, THSP2, ANG, and MUC16.STRA, achieved consistent performance in the validation set. The panel of CA19-9 with the glycan biomarker CA199.STRA improved sensitivity from 0.44 with 0.98 specificity for CA19-9 alone to 0.71 with 0.98 specificity (p < 0.001, 1000-fold bootstrap). Similarly, CA19-9 combined with the protein biomarker LRG1 and CA199.STRA improved specificity from 0.16 with 0.94 sensitivity for CA19-9 to 0.65 with 0.89 sensitivity (p < 0.001, 1000-fold bootstrap). We further validated significantly improved performance using biomarker panels that did not include CA19-9. This study establishes the effectiveness of a coordinated study of previously discovered biomarkers and identified panels of those biomarkers that significantly increased the sensitivity and specificity of early-stage PDAC detection in a rigorous validation trial.

## 1. Introduction

Pancreatic ductal adenocarcinoma (PDAC) is an extremely aggressive malignancy with an overall five-year survival rate of ∼13% in the United States and one of the worst prognoses of all cancers [1]. Approximately ∼15% of PDAC patients are identified when the tumor is confined to the pancreas and node-negative (localized stage) [1]. Remarkably, such patients have an ∼ 3-fold higher 5-year survival rate (44%) than patients with node positive (regional stage) PDAC (16%) [1,2], supporting the goal of pursuing strategies to identify PDAC patients when their disease is at an earlier, localized stage. This goal, however, has been constrained by the lack of a low-cost and convenient blood test that would perform well enough for surveillance of elevated-risk populations [3]. The best-known blood biomarker for PDAC, CA19-9, is elevated in the majority of PDAC patients but is not highly specific for PDAC, yielding 0.70-0.75 sensitivity and 0.70-0.75 specificity for distinguishing PDAC from non-cancer conditions [4,5]. Such performance is insufficient for surveillance for PDAC [3,6].

Previous studies have identified many candidate biomarkers for PDAC, although the best individual biomarkers identify PDAC patients with sensitivity and specificity no better than CA19-9 (see reviews [3,7,8]). Such biomarkers potentially identify separate subsets patients without a complementary identification of non-PDAC subjects, thus yielding superior performance if used in combination. However, the combined performance of independent, previously reported biomarkers is not known. Achieving the rigor necessary to ensure reproducibility is especially challenging in studies investigating multiple biomarkers or laboratories.

Here we investigated whether previously reported PDAC biomarkers used in combination as a panel could improve performance over CA19-9 alone in a rigorous validation study. To enable a rigorous validation study of biomarkers spanning multiple laboratories, we used a common set of specimens, consistent collection methods with appropriate inclusion/exclusion criteria, uniform sample handling and blinded distribution, and independent biostatistical analysis. The new panels were validated by applying classification rules with pre-determined cutoffs to an independent, blinded cohort. We assessed the reliability of the panels through the consistency between the training and validation sets, the identification of complementary subsets of patients without proportionally increasing false-positive detection, and improved performance over CA19-9 in combinations that did not include CA19-9. The coordinated study identified panels of biomarkers from separate laboratories that significantly improved specificity and sensitivity over CA19-9 and achieved the performance needed for surveillance among populations with an increased risk of PDAC.

## 2. Methods

### 2.1. Human Specimens

Human blood plasma specimens were assembled through the clinical centers at UPMC and MD Anderson. All specimens were collected by each site under the approval of their respective institutional review boards and were processed under a standard operating procedure approved by the Early Detection Research Network [4]. The samples were collected prior to any cancer treatment. Only de-identified information regarding the subjects was shared with the study researchers.

### 2.2. Study Design

The study biostatistician and the sites providing samples selected and coded the samples and assembled the metadata for a training and a validation set separately. The coded sample aliquots were sent to the individual laboratories for analysis. The training set cohort (**Table 1**) consisted of PDAC patients (n = 66) and control subjects (n = 66) that included the major, potentially confounding conditions in a surveillance setting: pancreatitis, benign cysts, and benign biliary obstruction. Most cases (80%) had resectable PDAC. The validation set cohort (**Table 2**) included patients with resectable PDAC (n = 170) and control subjects (n = 125) with a composition matching the training set.

**Table 1.** Demographics of PDAC cases and controls used in cross-laboratory panel development in the training set. The p values were calculated by chi-square tests for categorical variables and t-tests for continuous variables.

|  | Case (N=66) | Control (N=66) | p value | Total (N=132) |
| --- | --- | --- | --- | --- |
| **Site** |  |  |  |  |
| MD Anderson | 37 (56.1%) | 10 (15.2%) |  | 47 (35.6%) |
| Pittsburgh | 29 (43.9%) | 56 (84.8%) |  | 85 (64.4%) |
| **Gender** |  |  |  |  |
| Male | 35 (53.0%) | 24 (36.4%) | 0.05 | 59 (44.7%) |
| Female | 31 (47.0%) | 42 (63.6%) |  | 73 (55.3%) |
| **Race** |  |  |  |  |
| White | 62 (93.9%) | 64 (97.0%) | 0.11 | 126 (95.5%) |
| Black or African American | 0 (0%) | 2 (3.0%) |  | 2 (1.5%) |
| Asian | 1 (1.5%) | 0 (0%) |  | 1 (0.8%) |
| Other | 3 (4.5%) | 0 (0%) |  | 3 (2.3%) |
| **Hispanic** |  |  |  |  |
| Yes | 2 (3.0%) | 1 (1.5%) | 0.55 | 3 (2.3%) |
| No | 63 (95.5%) | 65 (98.5%) |  | 128 (97.0%) |
| Missing | 1 (1.5%) | 0 (0%) |  | 1 (0.8%) |
| **Age** |  |  |  |  |
| Mean (SD) | 65.9 (8.57) | 64.4 (10.9) | 0.40 | 65.1 (9.77) |
| Median (Min, Max) | 65.5 (41.0, 86.0) | 64.5 (25.0, 88.0) |  | 65.0 (25.0, 88.0) |
| **Smoking** |  |  |  |  |
| Never | 35 (53.0%) | 30 (45.5%) | 0.04 | 65 (49.2%) |
| Past Smoker | 22 (33.3%) | 15 (22.7%) |  | 37 (28.0%) |
| Current Smoker | 9 (13.6%) | 21 (31.8%) |  | 30 (22.7%) |
| **Diabetes** |  |  |  |  |
| No | 25 (37.9%) | 50 (75.8%) | <0.001 | 75 (56.8%) |
| Yes | 41 (62.1%) | 16 (24.2%) |  | 57 (43.2%) |
| **Type** |  |  |  |  |
| Resectable PDAC (stage I/II) | 53 (80%) | 0 (0%) |  | 66 (50.0%) |
| Nonresectable PDAC (stage III/IV) | 13 (20%) | 0 (0%) |  |  |
| MCN low grade | 0 (0%) | 2 (3.0%) |  | 2 (1.5%) |
| IPMN low grade | 0 (0%) | 11 (16.7%) |  | 11 (8.3%) |
| Benign cysts SCA pseudocysts retention | 0 (0%) | 6 (9.1%) |  | 6 (4.5%) |
| Pancreatitis | 0 (0%) | 15 (22.7%) |  | 15 (11.4%) |
| Benign Biliary obstruction | 0 (0%) | 8 (12.1%) |  | 8 (6.1%) |
| EUS with benign or normal pancreas | 0 (0%) | 19 (28.8%) |  | 19 (14.4%) |
| Healthy Control | 0 (0%) | 5 (7.6%) |  | 5 (3.8%) |

**Table 2.** Demographics of cases and controls used in cross-laboratory panel development in the validation set. The p values were calculated by chi-square tests for categorical variables and t-tests for continuous variables.

|  | Case (N=170) | Control (N=125) | p value | Total (N=295) |
| --- | --- | --- | --- | --- |
| **Site** |  |  |  |  |
| MD Anderson | 68 (40.0%) | 28 (22.4%) |  | 96 (32.5%) |
| Pittsburgh | 102 (60.0%) | 97 (77.6%) |  | 199 (67.5%) |
| **Gender** |  |  |  |  |
| Male | 86 (50.6%) | 52 (41.6%) | 0.13 | 138 (46.8%) |
| Female | 84 (49.4%) | 73 (58.4%) |  | 157 (53.2%) |
| **Race** |  |  |  |  |
| White | 153 (90.0%) | 115 (92.0%) | 0.13 | 268 (90.8%) |
| Black or African American | 10 (5.9%) | 7 (5.6%) |  | 17 (5.8%) |
| American Indian or Alaska Native | 1 (0.6%) | 0 (0%) |  | 1 (0.3%) |
| Asian | 0 (0%) | 2 (1.6%) |  | 2 (0.7%) |
| Other | 5 (2.9%) | 0 (0%) |  | 5 (1.7%) |
| Missing | 1 (0.6%) | 1 (0.8%) |  | 2 (0.7%) |
| **Hispanic** |  |  |  |  |
| No | 163 (95.9%) | 121 (96.8%) | 0.87 | 284 (96.3%) |
| Yes | 6 (3.5%) | 4 (3.2%) |  | 10 (3.4%) |
| Missing | 1 (0.6%) | 0 (0%) |  | 1 (0.3%) |
| **Age** |  |  |  |  |
| Mean (SD) | 65.4 (9.89) | 62.5 (13.7) | 0.03 | 64.2 (11.7) |
| Median (Min, Max) | 66.0 (35.0, 89.0) | 62.0 (18.0, 91.0) |  | 65.0 (18.0, 91.0) |
| **Smoke** |  |  |  |  |
| Never | 97 (57.1%) | 48 (38.4%) | 0.05 | 145 (49.2%) |
| Former | 51 (30.0%) | 44 (35.2%) |  | 95 (32.2%) |
| Current | 18 (10.6%) | 20 (16.0%) |  | 38 (12.9%) |
| Current Smokeless Tobacco | 1 (0.6%) | 0 (0%) |  | 1 (0.3%) |
| Former Smokeless Tobacco | 2 (1.2%) | 2 (1.6%) |  | 4 (1.4%) |
| Former Smoke and Smokeless | 1 (0.6%) | 4 (3.2%) |  | 5 (1.7%) |
| Missing | 0 (0%) | 7 (5.6%) |  | 7 (2.4%) |
| **Diabetes** |  |  |  |  |
| No | 126 (74.1%) | 91 (72.8%) | 0.62 | 217 (73.6%) |
| Yes | 43 (25.3%) | 27 (21.6%) |  | 70 (23.7%) |
| Missing | 1 (0.6%) | 7 (5.6%) |  | 8 (2.7%) |
| **Type** |  |  |  |  |
| Resectable PDAC | 170 (100%) | 0 (0%) |  | 170 (57.6%) |
| MCN low grade | 0 (0%) | 6 (4.8%) |  | 6 (2.0%) |
| IPMN low grade | 0 (0%) | 15 (12.0%) |  | 15 (5.1%) |
| Benign cysts | 0 (0%) | 16 (12.8%) |  | 16 (5.4%) |
| Pancreatitis | 0 (0%) | 30 (24.0%) |  | 30 (10.2%) |
| Benign biliary obstruction | 0 (0%) | 8 (6.4%) |  | 8 (2.7%) |
| EUS with benign or normal | 0 (0%) | 16 (12.8%) |  | 16 (5.4%) |
| Healthy control | 0 (0%) | 15 (12.0%) |  | 15 (5.1%) |
| Healthy control with family history | 0 (0%) | 19 (15.2%) |  | 19 (6.4%) |

We analyzed the resulting data in two ways. First, we optimized for specificity while controlling for high (∼0.95) sensitivity, and second, we optimized for sensitivity while controlling for high (∼0.95) specificity. Using the data from the training set, we developed classification rules and determined corresponding cutoffs for making case/control calls. We then applied the classification rules with fixed cutoffs to the validation set without further optimization to determine the statistical significance and potential future performance of the biomarker panels. Using CA19-9 as a benchmark, we tested the statistical significance of improvement over CA19-9 using the dichotomized case/control calls in the blinded validation set.

### 2.3. Biomarker Assays

The clinical-assay CA19-9 measurements were obtained from UPMC’s diagnostics laboratory using the GI Monitor immunoenzymatic assay on the Beckman Coulter DXI 800 instrument. The assay results were obtained either from pre-treatment testing as part of routine medical care within two weeks of each subject’s sample collection or for research purposes specific to this study.

The biomarker assays were run at the contributing laboratories according to the protocols used in the initial validation of the biomarkers. The assays at VAI used antibody microarrays with fluorescence detection [9]. The assays at MD Anderson and UPMC used Luminex bead-based immunoassays using Milliplex (MilliporeSigma) kits according to the provided protocol [10,11]. The assays at UNMC used a previously described sandwich ELISA assay [12]. See Supplementary Information for additional details.

### 2.4. Statistical Analyses

#### Performance Estimation of Individual Candidate Markers

The performance of individual candidate markers and CA19-9 for separating PDAC from benign and health controls in the training and validation set was evaluated using the receiver operating characteristics (ROC) curve and area under the ROC curve (AUC) (for markers measured on the continuous scale) and using sensitivity and specificity (when there is an established threshold for making binary case/control calls). Nonparametric bootstrap percentile confidence intervals were generated for performance estimates with 1,000 bootstrap samples stratified on case/control status.

#### Biomarker Panel Development and Validation

Cross-lab biomarker panels were developed using the training set samples, targeting either high specificity or high sensitivity and improvement over clinical CA19-9. Parsimonious panels of 2-4 biomarkers were developed by adding new markers to CA19-9, based on least absolute shrinkage and selection operator (LASSO), random forest, and Logic OR rule, with best panels of different sizes selected based on cross-validated classification performance. In particular, i) the training set was randomly split into 2:1 fold stratified on case/control status to serve as the training and test subset, respectively; a biomarker panel (and corresponding binary rule with threshold chosen for 95% specificity or 95% sensitivity) was developed from the training subset, performance (sensitivity and specificity) of which is then estimated from the test subset; this random-split process was repeated 1,000 times with average test performance computed. ii) the algorithms resulting in panels of top performance (i.e., sensitivity at high specificity or specificity at high sensitivity) with 2-4 biomarkers were then applied to the full training set for panel development and cutoff estimation. iii) performance of estimated panels from training set was evaluated in validation set in terms of sensitivity and specificity at the pre-specified binary panel call. The incremental value of the panel over CA19-9 alone was assessed based by the average of sensitivity and specificity for the binary panel call and the binary CA19-9 call. P values were determined by standard error estimated from 1000 bootstrap samples. An analogous strategy was used to develop and validate cross-lab panels that do not include CA19-9.

## 3. Results

### 3.1. Biomarkers with Complementary Value to CA19-9

Four independent laboratories performed assays on previously discovered protein, glycan, and protein-glycoform pancreatic cancer biomarkers (**Fig. 1 and Table 3**). We used CA19-9 values from a clinical assay performed by the UPMC Diagnostics Laboratory and two additional CA19-9 assays in the laboratories at UPMC and VAI. The UPMC and VAI assays showed inter-assay correlations with the clinical CA19-9 assay of 0.90-0.91 across all samples and had nearly equivalent area-under-the-curve values in ROC analysis for discriminating the cancer cases from controls (**Supplementary Fig. 1**). We thus used the clinical CA19-9 values in the subsequent analyses; similar results were observed using the lab-determined CA19-9 values (not shown).

**Table 3.** Biomarkers tested in panel development. UPMC, University of Pittsburgh Medical Center; VAI, Van Andel Institute; MCACC, MD Anderson Cancer Center; UNMC, University of Nebraska Medical Center.

| Site | Biomarker | Type | Method | Reference |
| --- | --- | --- | --- | --- |
| UPMC | TGM2 | Protein | Luminex |  |
|  | THSP2 | Protein | Luminex |  |
|  | HEP | Protein | Luminex |  |
|  | TIMP2 | Protein | Luminex |  |
|  | ANG | Protein | Luminex |  |
| VAI | CA199.STRA | Glycan | Antibody array | [9] |
|  | MUC5AC.STRA | Protein glycoform | Antibody array | [9] |
|  | MUC16.STRA | Protein glycoform | Antibody array | [9] |
| MDACC | CA19-9 | Protein | Luminex | [10,11] |
|  | TIMP1 | Protein | Luminex | [10,11] |
|  | LRG1 | Protein | Luminex | [10,11] |
| UNMC | MUC4 | Protein | ELISA | [12] |
|  | MUC5AC | Protein | ELISA | [12] |
| UPMC Diagnostics Laboratory | CA19-9 | Glycan | Clinical assay | NA |

**Figure 1.**
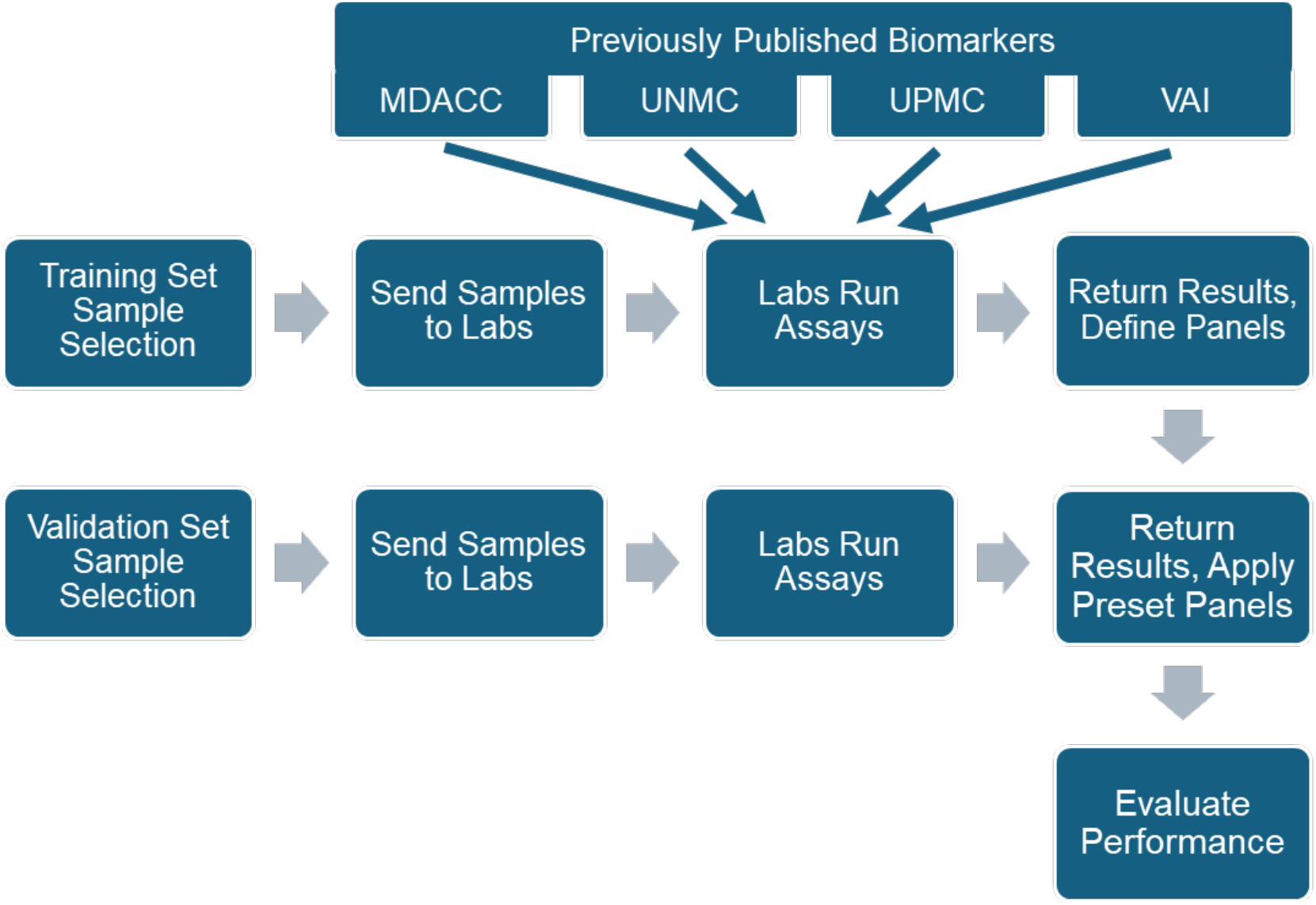
Flowchart of the Study Design.

The CA19-9 biomarker provided a benchmark. At the predetermined, clinically relevant cutoff of 37 U/mL, CA19-9 gave 0.76 sensitivity and 0.79 specificity (**Supplementary Table 1**). At a cutoff optimized for high specificity (164 U/mL), CA19-9 gave 0.50 sensitivity and 0.95 specificity, and at a cutoff optimized for high sensitivity (2.0 U/mL), CA19-9 gave 0.95 sensitivity and 0.02 specificity (**Supplementary Tables 1 and 2**). The low cutoff necessary to give 0.95 sensitivity is not relevant for distinguishing varying levels of CA19-9 (as indicated by the very low specificity), highlighting the inability of CA19-9 to detect PDAC with high sensitivity.

We next evaluated the performance of panels of CA19-9 with additional biomarkers. Given the large number of potential panels of assays, the algorithm selected parsimonious panels of biomarkers (2-4 biomarkers) with the best performance in the cross-validation analysis (see Methods). Several panels of biomarkers that included CA19-9 improved cross-validated performance over CA19-9 in the training data (**Table 4 and Fig. 2A**). In the high-specificity analysis, the panel of CA19-9 with CA199.STRA (a sandwich immunoassay using capture of the CA19-9 glycan and detection of the STRA glycan [9]) gave 0.63 sensitivity and 0.92 specificity. The CA199.STRA marker also improved upon CA19-9 as an individual marker (**Supplementary Table 1 and Supplementary Fig. 2**). In the high-sensitivity analysis, several 2-marker and 3-marker panels improved over CA19-9 alone (**Table 4 and Fig. 2B**). Thus, several biomarker combinations substantially improved both sensitivity and specificity relative to CA19-9 alone. The classification rules and cutoffs defined from the training set (**Supplementary Table 2**) were used in all subsequent analyses.

**Table 4.** Biomarker performance in the training set. Avg, average of sensitivity and specificity based on the cross-validated performance. Most of the panels were derived from a binary OR rule (Supplementary Table 2), not a continuous score, for which the calculation of area-under-the-curve in receiver-operator-characteristic analysis is not relevant.

|  |  | Basis Biomarker | Additional Biomarkers | Sensitivity | Specificity | Avg | Method |
| --- | --- | --- | --- | --- | --- | --- | --- |
| With CA19-9 | High specificity | Clinical CA19-9 | - | 0.49 | 0.94 | 0.71 | - |
|  |  | Clinical CA19-9 | CA199.STRA | 0.63 | 0.92 | 0.77 | OR |
|  |  | Clinical CA19-9 | CA199.STRA + TIMP1 | 0.66 | 0.96 | 0.81 | OR |
|  |  | Clinical CA19-9 | CA199.STRA + ANG | 0.65 | 0.95 | 0.80 | OR |
|  |  | Clinical CA19-9 | CA199.STRA + MUC16.STRA | 0.67 | 0.95 | 0.81 | OR |
|  |  | Clinical CA19-9 | CA199.STRA + TGM2 + HEP | 0.63 | 0.95 | 0.79 | RF |
|  | High sensitivity | Clinical CA19-9 | - | 0.95 | 0.06 | 0.51 | - |
|  |  | Clinical CA19-9 | TIMP1 | 0.92 | 0.50 | 0.71 | OR |
|  |  | Clinical CA19-9 | LRG1 | 0.90 | 0.49 | 0.70 | OR |
|  |  | Clinical CA19-9 | THSP2 | 0.90 | 0.48 | 0.69 | OR |
|  |  | Clinical CA19-9 | CA199.STRA + TIMP1 | 0.95 | 0.65 | 0.80 | OR |
|  |  | Clinical CA19-9 | CA199.STRA + LRG1 | 0.95 | 0.63 | 0.79 | OR |
|  |  | Clinical CA19-9 | CA199.STRA + LRG1 + MUC5AC | 0.91 | 0.44 | 0.68 | Logi-Reg |
|  |  | Clinical CA19-9 | CA199.STRA + THSP2 + TIMP1 | 0.94 | 0.36 | 0.65 | RF |
| No CA19-9 | High specificity | CA199.STRA | ANG | 0.59 | 0.92 | 0.75 | OR |
|  |  | CA199.STRA | MUC16.STRA | 0.62 | 0.92 | 0.77 | OR |
|  |  | CA199.STRA | MUC5AC | 0.64 | 0.93 | 0.79 | Logi-Reg |
|  |  | CA199.STRA | THSP2 + MUC5AC | 0.48 | 0.94 | 0.71 | RF |
|  |  | CA199.STRA | THSP2 + MUC5AC + ANG | 0.57 | 0.93 | 0.75 | Logi-Reg |
|  | High sensitivity | CA199.STRA | MUC16.STRA | 0.91 | 0.47 | 0.69 | OR |
|  |  | CA199.STRA | LRG1 | 0.90 | 0.53 | 0.71 | OR |
|  |  | CA199.STRA | TIMP1 | 0.91 | 0.49 | 0.70 | OR |
|  |  | CA199.STRA | THSP2 + MUC5AC | 0.94 | 0.23 | 0.58 | RF |
|  |  | CA199.STRA | THSP2 + MUC5AC + ANG | 0.91 | 0.44 | 0.67 | Logi-Reg |
|  |  | CA199.STRA | THSP2 + MUC5AC + TIMP1 | 0.95 | 0.42 | 0.69 | RF |

**Figure 2.**
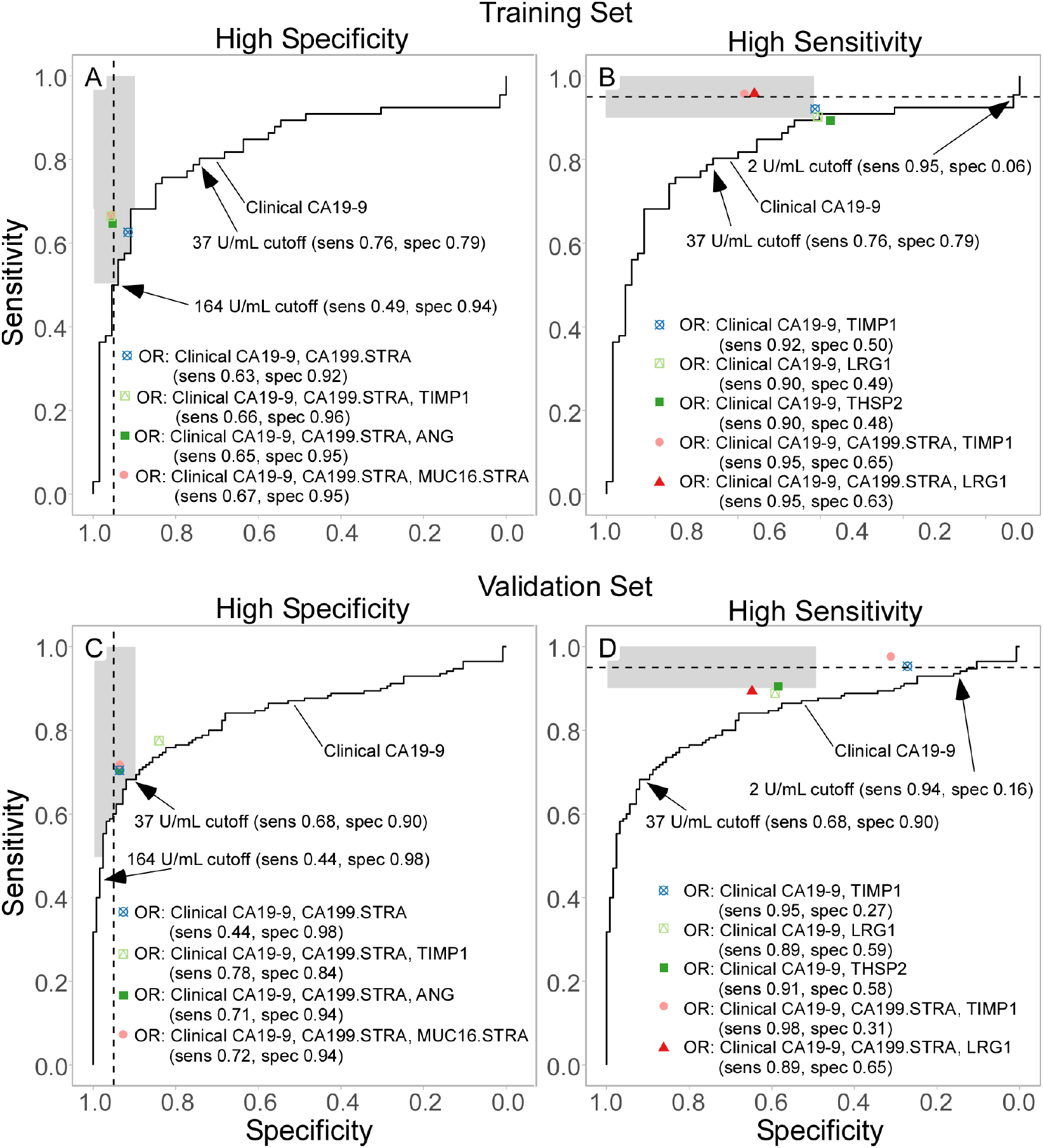
Training Set and Validation Set Performance. The receiver-operator characteristic curves for CA19-9 are shown with the point values of the biomarkers panels developed for A) high specificity and B) high sensitivity in the training set (with cross-validation) and for C) high specificity and D) high sensitivity in the validation set. The gray boxes indicated the target regions of performance enhancement.

### 3.2. Validation of Biomarker Panels Yielding Improved Specificity or Sensitivity

Using the pre-determined classification rules and cutoffs in the validation set, CA19-9 by itself had performance that was insufficient for surveillance: 0.68 sensitivity and 0.90 specificity using the 37 U/mL cutoff (**Supplementary Table 3**); 0.44 sensitivity and 0.98 specificity using the high-specificity cutoff (164 U/mL); and 0.94 sensitivity and 0.16 specificity using the high-sensitivity cutoff (2 U/mL) (**Table 5 and Fig. 2D**). Among the individual biomarkers, CA199.STRA exhibited highest sensitivity (0.59 at 95% sensitivity) and MUC5AC exhibited best specificity (0.23 at 95% sensitivity) (**Supplementary Table 3**).

**Table 5.**
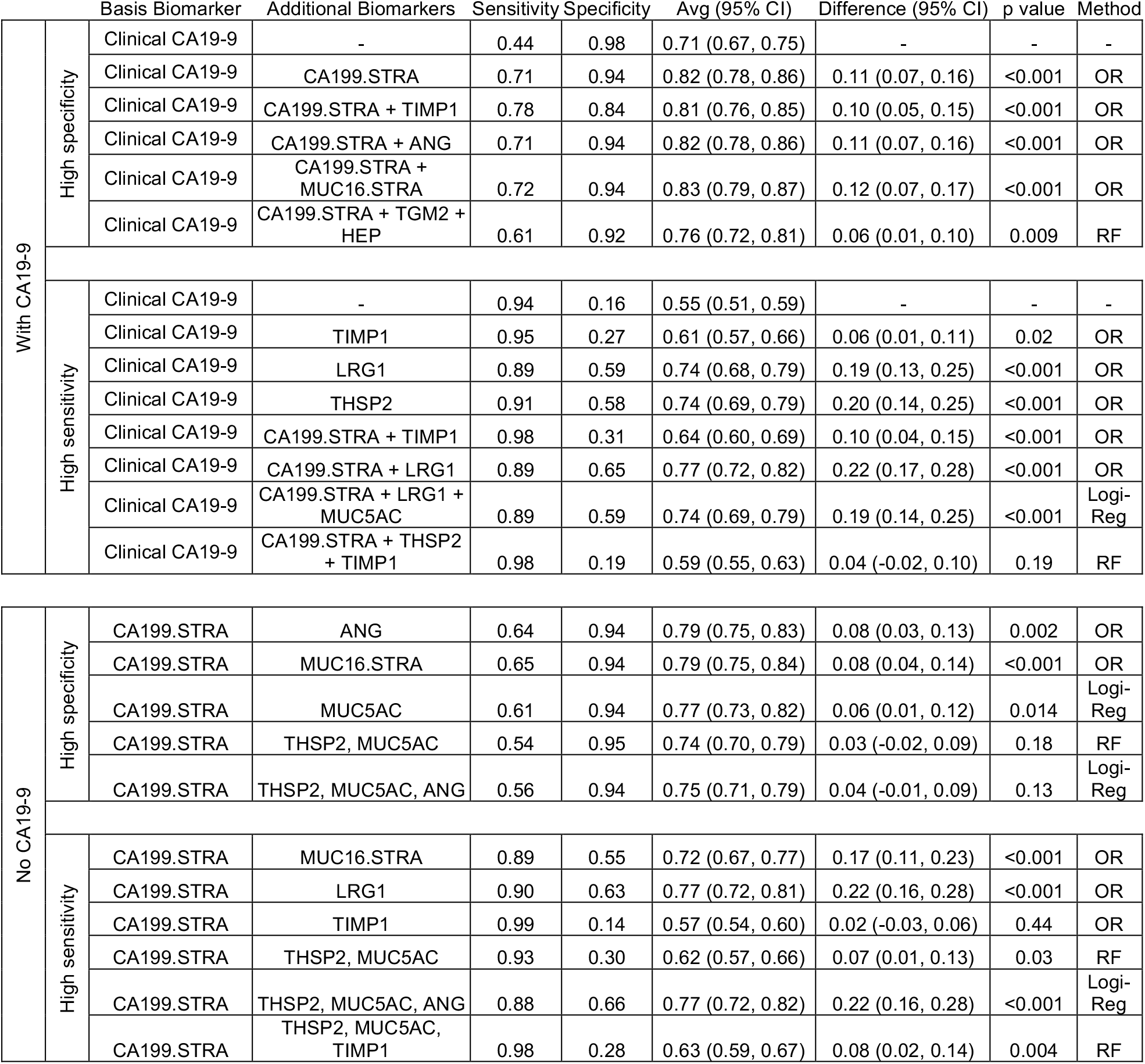
Biomarker performance in the validation set. Avg, average of sensitivity and specificity; CI, confidence interval; Difference, difference between CA19-9 and each panel (i.e. performance of biomarker panel minus the performance of CA 19-9) in the average of sensitivity and specificity.

The majority of biomarker panels from the training set demonstrated consistent performance in the validation set (**Table 5 and Fig. 2C**). The top-performing panel in the high-specificity analysis was CA19-9 and CA199.STRA. The average of sensitivity and specificity was significantly improved from 0.71 (95% C.I. 0.67-0.75) for CA19-9 alone to 0.82 (95% C.I. 0.78-0.86) (p < 0.001, standard error estimated from 1000-fold bootstrap samples). The top 2-marker panel in the high-sensitivity analysis, CA19-9 and THSP2, significantly improved the average sensitivity and specificity over CA19-9 to 0.74 (0.69, 0.79) (p < 0.001, 1000-fold bootstrap analysis). The 3-marker panel of CA19-9, CA199.STRA and LRG1 also significantly improved performance, to 0.77 (0.72, 0.82) (p < 0.001, 1000-fold bootstrap analysis), and was highly consistent with the results from the training set (**Compare Fig. 2B to Fig. 2D**). These data demonstrate reproducible, significant improvement upon CA19-9 alone for PDAC detection using pre-determined classification rules and cutoffs in blinded samples.

### 3.3. Complementary Subset Detection

We further tested the robustness of the improved sensitivity and specificity of the new biomarker panels by examining the extent of overlap in the patients and controls identified by CA19-9 alone or by the biomarker panel of CA19-9 and CA199.STRA. In the high-specificity analysis of the validation set, the panel identified about half of the cases not identified by CA19-9 alone at either the 37 U/mL or the 164 U/mL cutoff (23/54 (43%) and 50/96 (52%) cases, respectively), whereas CA19-9 alone at the respective cutoffs identified 38% (19/50) and 8% (4/50) of the cases not identified by the panel (**Figs. 3A and 3B**). The new panel had a low false-positive detection rate of controls not detected by CA19-9 alone: 5.3% (6/112) and 5.7% (7/123) at the respective cutoffs.

**Figure 3.**
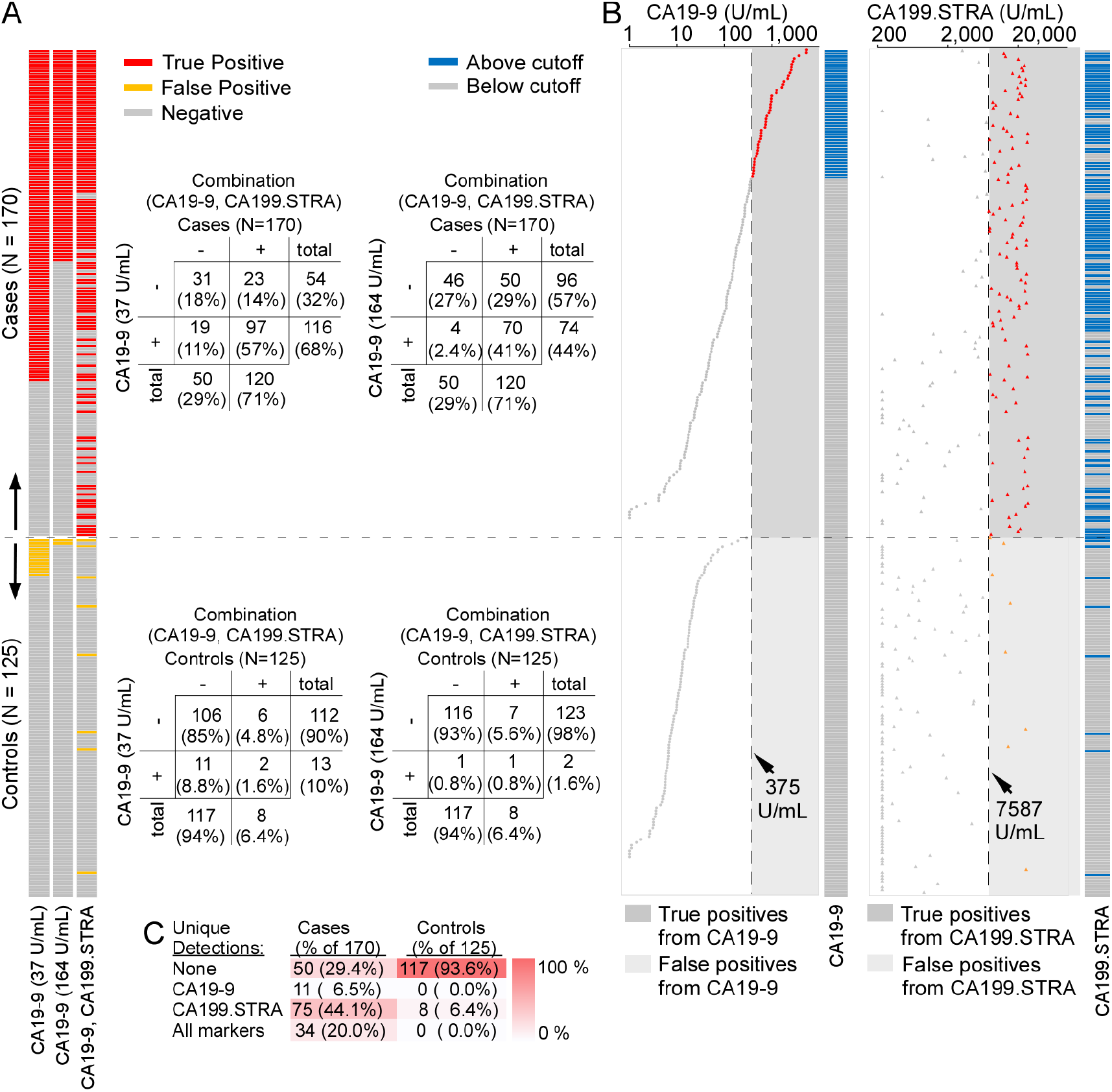
Complementary Enhancement of Sensitivity with High Specificity. A) Biomarker-based classifications of the cases and controls in the validation set, targeting high specificity. The cases and controls are ordered by descending CA19-9 values. B) CA19-9 (left) and CA199.STRA (right) values, ordered by descending CA19-9. The vertical dashed line indicates the cutoff used for each biomarker. The individual biomarker high/low status and the classification based on the panel are shown at right. If a specimen was elevated in either biomarker, it was classified as a case. C) Unique and common elevations in the two biomarkers in the cases and controls.

We examined whether individual samples were variously detected by only CA19-9 or only CA199.STRA in the biomarker panel. Of the 120 cases identified by either CA19-9 or CA199.STRA, 75 (44%) were uniquely identified by CA199.STRA, 11 (6.5%) uniquely by CA19-9, and 34 (20%) by both, and of the 8 controls misidentified as cases by the new panel, all 8 (6.4%) were due to the CA199.STRA assay (**Fig. 3C**). Thus, the improvement in sensitivity results from the low overlap and high unique detection by CA19-9 and CA199.STRA in the cases above the biomarkers cutoffs.

The most sensitive biomarker panel—CA19-9, CA199.STRA, and LRG1—significantly reduced the number of false-positive identifications while identifying nearly all cases (**Fig. 4A and 4B**), as did several other cross-laboratory panels. This new panel identified nearly all the cases identified by CA19-9 alone at either the 37 U/mL or the 2 U/mL cutoff, but it greatly reduced the number of false-positive identifications of controls relative to CA19-9. CA19-9 at 2 U/mL gave 82 false positive detections (66% of the 125 controls) that were not falsely identified by the panel, but the panel only gave 4 false positive detections (3.2% of the 125 controls) that were not falsely detected by CA19-9 (**Fig. 4A**). Using the predetermined cutoffs (**Supplementary Table 2**) for each biomarker, each assay in the new panel identified unique sets of patients (**Fig. 4B**). Each component of the new biomarker panel uniquely identified 0.6-35% of the 170 cases but only had 0-18% false positive detections out of the 125 controls (**Fig. 4C**). Therefore, the improved specificity of the new panel of CA19-9, CA199.STRA, and LRG1 over CA19-9 alone is a result of the minimal contribution of false positives from each member of the panel, along with complementary true-positive identification from each individual biomarker assay.

**Figure 4.**
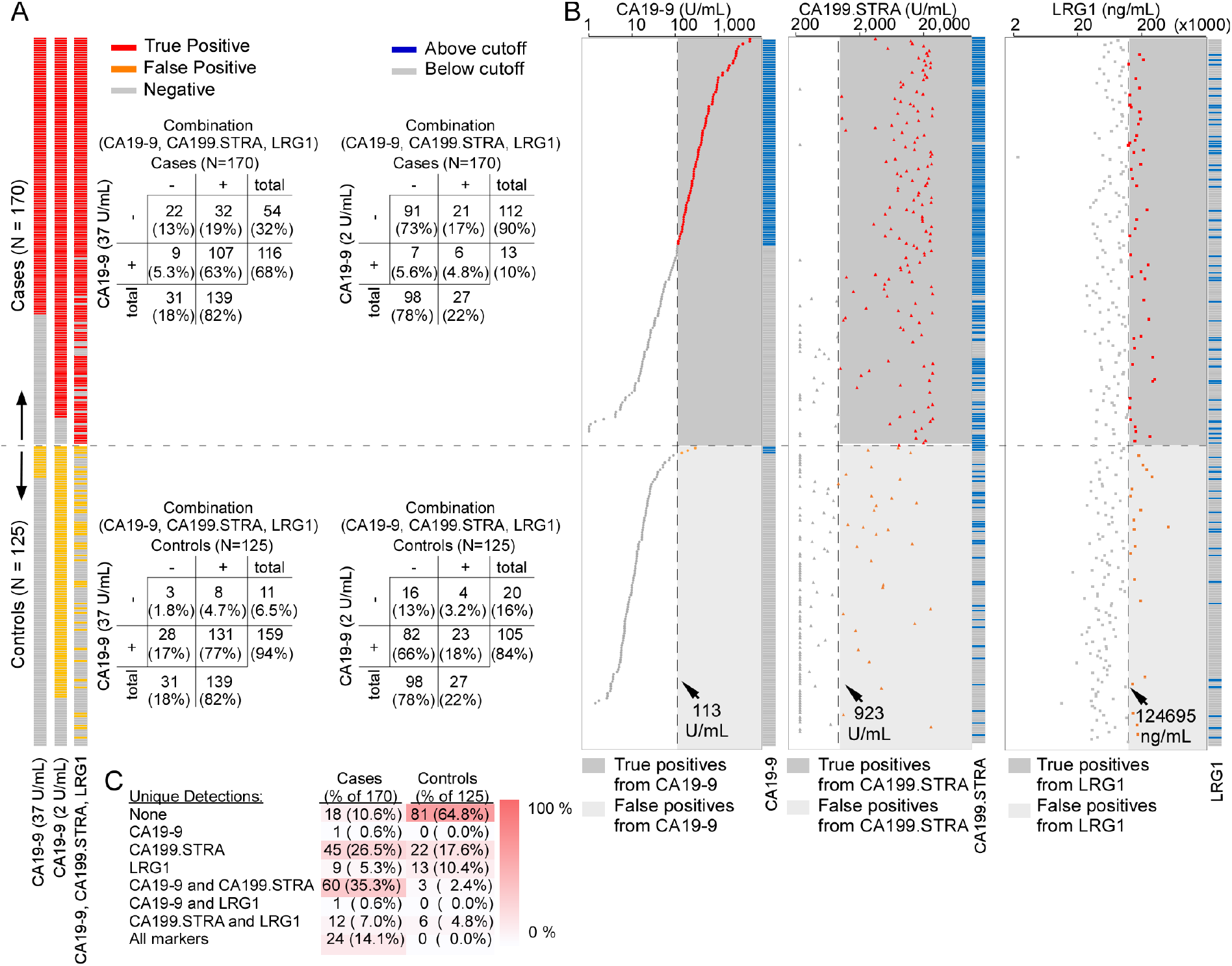
Complementary Enhancement of Specificity with High Sensitivity. A) Biomarker-based classifications of the cases and controls in the validation set, targeting high sensitivity. The cases and controls are ordered by descending CA19-9 values. B) CA19-9 (left), CA199.STRA (middle) and LRG1 (right) values, ordered by descending CA19-9. The vertical dashed line indicates the cutoff used for each biomarker. The individual biomarker’s high/low status and the classification based on the panel are shown at right. If a specimen was elevated in any biomarker, it was classified as a case. C) Unique and common elevations in the three biomarkers in the cases and controls.

### 3.4. Improved Performance Independent of CA19-9

A further test of the improved performance of the biomarkers is whether new biomarker panels that did not include CA19-9 also improved upon CA19-9 alone. In the high-specificity analysis of the training set, CA199.STRA alone (**Supplementary Table 1**) or in other new biomarker panels performed as well as any panel that included CA19-9 (**Table 4**). In the training set high-sensitivity analysis, new panels including CA199.STRA, THSP2, MUC5AC, and ANG or TIMP1 significantly improved upon CA19-9 alone and achieved performance comparable to new panels that included CA19-9. The improvements were consistent between the training and validation sets (**Supplementary Fig. 3**). In the validation set, CA199.STRA alone (**Supplementary Table 2**) or in a new panel with MUC5AC, MUC16.STRA, or ANG, achieved 0.61-0.65 sensitivity and 0.94 specificity, representing a significant improvement over CA19-9 alone (p < 0.001, 1000-fold bootstrap analysis) (**Table 5**). In the high-sensitivity analysis of the validation set, the new biomarker panel of CA199.STRA and LRG1 performed better than any other, significantly improving upon CA19-9 (p = 0.0004, 1000-fold bootstrap analysis) with 0.90 sensitivity and 0.63 specificity. Thus, new biomarkers panels that did not include CA19-9 performed better than CA19-9 alone.

## 4. Discussion

Here, for the first time, we report a rigorous study of previously discovered PDAC biomarkers used in combination to specifically and sensitively identify PDAC. While many previous studies have identified independent PDAC markers, a coordinated study of the combined value of the known PDAC biomarkers has not been demonstrated. In a multi-laboratory validation study, we identified increased sensitivity and specificity of new panels of known PDAC biomarkers over CA19-9 alone.

The robust, consistent performance of the panels in the training and validation sets likely stems from the use of previously discovered biomarkers. The CA199.STRA biomarker was discovered in 2016 in a study of glycans that are structurally related to CA19-9 and that could have complementary value for detection [13], and it was subsequently shown to have a value as a tissue and serological biomarker for pancreatic cancer [9]. The LRG1 protein was identified as a candidate biomarker through proteomics analysis of colorectal cancer cell lines [14] followed by enzyme-linked immunosorbent assay analysis of serum from pancreatic cancer patients [10,15]. The MUC5AC protein has long been recognized as a biomarker that is present in pancreatic cancer tissue through studies using monoclonal antibodies, and more recent studies demonstrated the value of MUC5AC as a blood biomarker for pancreatic cancers [12]. Therefore, all biomarkers investigated already had initial validation as PDAC-relevant biomarkers, providing a good foundation for yielding reliable results in rigorous validation.

The validation of the new biomarker panels was assessed by applying classification rules with cutoffs determined in the training data to the validation set, a process overseen by an independent biostatistician. This approach is rigorous and stands in contrast to the methods used in many previous studies that rederived cutoffs or classification rules in validation sample sets. By adhering strictly to established cutoffs and classification rules, we ensured a statistically robust and unbiased validation of performance [16]. The high consistency observed between the training and validation sets confirms that our panels of biomarkers were not overfitted. In other words, they were not tailored to the training set data in a post-hoc manner and remain relevant to future datasets. The study design and analysis thus fulfilled the principles of predictability, computability, and stability described in a previous report [17].

The validation was also demonstrated by the ability of biomarkers to identify different PDAC subsets without increasing the number of false positive cases identified. This feature is important for identifying PDACs across the heterogeneous groups of cancer cell subpopulations. Gene-expression profiling studies on whole-tumor mRNA have uncovered two main PDAC-cell subtypes, classical and basal (or squamous) (4–7). The two main subtypes, along with other, less-well characterized PDAC cancer cell subpopulations [18], typically coexist within tumors in greatly varying proportions. This molecular variation could explain why our new biomarker panels detect more cancers than a lone biomarker assay. Studies linking tumor characteristics to peripheral blood levels will help clarify such associations.

In addition, the validation of the new biomarker panels was established by the improved performance over CA19-9 even when CA19-9 was not included in the panel. For example, the 2-biomarker panel of CA199.STRA and LRG1 yielded 0.90 sensitivity and 0.63 specificity compared to 0.94 sensitivity and 0.16 specificity for CA19-9 alone (**Table 5**), a significant improvement. This benchmark stands in contrast to previous studies that have focused on demonstrating incremental improvement in combination with CA19-9.

The performance achieved here to identify more PDAC patients when the disease is still in an early stage warrants further development. Surveillance by radiographic imaging via magnetic resonance imaging (MRI), computed tomography (CT), or endoscopic ultrasound (EUS) [7] is already practiced in some centers among individuals with a strong family history of PDAC or a panel of family history and a known germline pathogenic variant associated with risk [19]. Recent studies have indicated that surveillance among such individuals increased the percentage of PDAC patients diagnosed with early-stage disease with concomitantly improved outcomes [20–22]. Among additional elevated-risk groups, such as those with chronic pancreatitis, sudden-onset diabetes [19], or PDAC-associated clinical histories [23], surveillance by radiographic imaging would not be sustainable owing to a lower prevalence of PDAC [3,7]. However, a blood test could be a cost-effective way to identify patients who could be further evaluated by radiographic imaging. Such a biomarker would need to be sensitive and specific enough to enrich the prevalence of PDAC to the point where follow-up imaging would be sustainable. As an example, a biomarker with 0.95 specificity and 0.65 sensitivity—as achieved here—used to screen for PDAC among patients with sudden-onset diabetes, among whom the prevalence of early-stage PDAC is estimated to be 0.8% [24,25], would enrich the prevalence of PDAC to 9.5% [6]. This prevalence is higher than the PDAC prevalence among individuals with familial PDAC, estimated to be 1.6% [20], and therefore easily within the range warranting follow-up by imaging.

The current study had several limitations that should be addressed in future research. The study did not use consecutive, prospectively collected and analyzed samples, which would better assess clinical value than banked, retrospective samples. The number of samples analyzed, though providing high statistical significance, needs to be increased to confirm ongoing, general value for future applications fully. In addition, the samples were collected around the time of diagnosis of early-stage PDAC rather than from a surveillance study. This design was adopted because samples collected in surveillance are available only in limited quantities that are prohibitive for use in a multi-laboratory study. We designed the current study to minimize the effects of such limitations, but the full assessment of the biomarker panels validated here will require prospective studies using a clinical assay in a surveillance setting.

It will be important in future studies to investigate each of the biomarkers for specific uses. For example, MUC5AC previously showed improvements over CA19-9 for the detection of resectable PDAC relative to non-resectable PDAC [12]. This is consistent with its better performance in the validation set (AUC 0.71; sensitivity 0.25 and specificity 0.23), which included only resectable PDAC, as compared to the training set (AUC 0.53; sensitivity 0.02 and specificity 0.00) (**Supplementary Tables 1 and 3**), which included both resectable and non-resectable PDAC. Thus, the improved performance of MUC5AC corroborates our observations where MUC5AC performed better than CA19.9 in identifying resectable PDAC cases [12]. Likewise, LRG1, ANG, THSP2 showed promise for settings where high sensitivity is preferred, as needed to rule out PDAC rather than to rule out non-PDAC. These observations also make a case for evaluating the performance of each of the biomarkers as an anchor and building a panel of combination markers with and without CA19-9. The further mining of the full dataset together with future panel studies, could reveal such value.

In summary, the new panels of previously discovered biomarkers significantly increased the sensitivity and specificity of early-stage PDAC detection, demonstrating the future feasibility of surveillance for PDAC using panels of serum assays. Studies are ongoing to determine the performance of these panels in samples collected prior to the diagnosis of PDAC.

## Acknowledgements

Funding was received from U01CA200466, U01CA200468, U01CA152653, U01CA226158, U24CA086368 and Sheikh Khalifa bin Zayed Foundation. David M Brass, PhD, assisted in the preparation of this manuscript.

