## Supplementary Information for "A Rigorous Multi-Laboratory Study of Known PDAC Biomarkers Identifies Increased Sensitivity and Specificity Over CA19-9 Alone"

### Contents

- Supplementary Methods.
  - Biomarker Assays
  - Statistical Analyses
- Supplementary Table 1. Individual Biomarker Performance in the Training Set.
- Supplementary Table 2. Cutoffs and Classification Rules of Biomarker Panels.
- Supplementary Table 3. Individual Biomarker Performance in the Validation Set.
- Supplementary Figure 1. Comparison of Clinical CA19-9 with the Two Research CA19-9 Results.
- Supplementary Figure 2. Receiver-Operator Characteristic (ROC) Curves from The Training Set for Each Lab's Individual Biomarkers Compared with Clinical CA19-9.
- Supplementary Figure 3. Training Set and Validation Set Performance of Combinations without CA19-9.

### Supplementary Methods

#### Biomarker Assays

*Van Andel Institute*. The assays were run using antibody microarrays with fluorescence detection [8]. The capture antibodies (CA19-9 (clone 1116-NS-19-9), MUC5AC (clone 45M1), and MUC16 (clone X325)) were prepared at a concentration of 250 µg/mL in a buffer of phosphate buffered saline (PBS) with 15% glycerol and 0.005% Tween-20. The antibody solutions were deposited onto nitrocellulose covered microscope slides (PATH slides, Grace BioLabs) in arrays using the Aushon 2470 microarrayer. Each antibody was printed in 3 replicate spots in arrays of 4 x 4 with a spacing of 280 microns between spots. Each microscope slide contained 96 replicate arrays in a checkerboard 8 x 24 grid with 4.5 mm spacing. The arrays were segregated by wax borders (Slide Imprinter, The Gel Company) and blocked for one hour at RT with 1% BSA in PBS with 0.5% Tween-20. Plasma samples were diluted 1:8 and 1:32 (v:v) in sample buffer (Tween-20, Brij-35, PBS, protease inhibitor (Complete Mini, EDTA-free), IgG cocktail (ChromPure Mouse IgG, 200 µg/mL)) and incubated on the arrays in duplicate wells for each dilution overnight at 4 °C. Samples used for TRA (clone TRA-1-60) detection were then incubated with α2-3 neuraminidase S (40 U/mL) at 37 °C for one hour. Biotinylated primary detection antibodies (CA19-9 and TRA-1-60) were applied at 3 µg/mL in detection buffer (PBS, 0.1% BSA, 0.05% Tween-20) for one hour at RT, followed by streptavidin-conjugated Cy5 (2 µg/mL in the same detection buffer) also for one hour at RT.

A fluorescence scanner detected a fluorescent signal (InnoScan 1100 AL, Innopsys). The signal was quantified using SignalFinder-MA [31] and calibrated using standards randomized throughout the slides. Standard curves were generated using 11 serial twofold dilutions of a CA19-9 standard (CA19-9 Calibrator Grade, Ray Biotech). The calibration of the experimental data was performed using custom software written in Matlab. The calibrated values were corrected for dilution and averaged across replicates. Each sample was run in three independent experiments, and the results were averaged across experiments.

*MD Anderson.* Plasma protein concentrations for CA19-9, LRG1 and TIMP1 were determined via Luminex bead-based immunoassays using Milliplex (MilliporeSigma) kits HCCBP1-58MAG (CA19-9), HCVD6MAG-67K (LRG1) and HTMP1MAG-54K (TMP1) according to the provided kit protocol as previously described [29,32]. The plasma samples with internal controls were thawed overnight at 4°C and diluted using kit diluent (1:6 for CA19-9, 1:2000 for LRG1, and 1:50 for TIMP1). Assay plates comprised singlet assay samples and duplicate 8-point standard dilution series and high/low QCs for each protein marker. Milliplex antibody-immobilized magnetic capture beads were added to the plates and incubated 18 hours overnight at 4°C with shaking at 750 RPM. The following day, the sample plates were 3x washed using a BioTek405TS plate washer. Milliplex biotinylated detection antibodies were added and incubated for 1 hour at RT with shaking at 750 RPM. Streptavidin/R-Phycoerythrin conjugate (SAPE) was added to the plate wells for 30 minutes at RT with shaking, followed by 3x washing and aspiration. Luminex Drive Fluid was added to the samples, and the plates were shaken for 5 minutes prior to reading on a Luminex MAGPIX instrument. The resulting assay data was analyzed with Milliplex Analyst software to yield sample mean fluorescence intensity and analyte concentration according to 5PL (5-parameter logistic) curve fitting of the respective assay standards.

*UPMC.* The proteins TGM2, THSP2, HEP, TIMP2, and ANG were analyzed using the Luminex bead-based immunoassays using Milliplex (MilliporeSigma) kits HCCB4MAG-58K (TGM2 and HEP), HANG2MAG-12K (THSP2 and ANG), and LKT003 (TIMP2). All steps followed the manufacturer's protocol as described above. The plasma samples were diluted into kit diluent (1:5 for TGM2, THSP2, HEP, and ANG) or into calibrator diluent RD6-48 (1:10 for TIMP2). All proteins were analyzed with the Bio-Plex 200 reader.

*UNMC.* MUC5AC levels were measured using a previously described sandwich ELISA assay [30]. 96-well microplates were coated with 10 μg/mL of capture antibody (mouse anti-MUC5AC, clone 13M1) diluted in 0.05 M carbonate buffer and incubated overnight at RT. The following day, the plates were washed twice with PBS with 0.1% Tween-20 (PBS-T) and blocked with 3% filtered BSA for 2.5–3 h at 37 °C, followed by three washes with PBS-T. Sample or standards (one-time prepared lysate of A549 cells expressing high levels of MUC5AC), appropriately diluted in 1% BSA, were added to each well. Plates were sealed and incubated overnight at 4 °C. Plates were washed four times with 0.1% PBS-T. Biotinylated mouse anti-MUC5AC mAb-1 (clone 45M1, Neo-Markers, Fremont, CA), diluted in 1% BSA, was added to each well. Plates were incubated for 2 h at 37°C. After washing, streptavidin poly-HRP, diluted to 0.0002 mg/mL in 1% BSA, was added to each well and incubated for 30 minutes at RT in the dark. Following washing, 100 μL of substrate (3,3′,5,5′-tetramethylbenzidine (TMB)) solution was added to each well, and the plates were incubated for 20 min at RT in the dark. The reaction was quenched by the addition of 1 M sulfuric acid, and the absorbance at 450 and 650 nm was measured using a microplate reader. The levels of MUC5AC were quantified as nanograms per mL of total protein in the A549 cell line of lung cancer, and the values were natural-log transformed. Values that were 0 before log transformation were replaced by half the minimum positive value of the marker.

**Statistical Analyses**

*Comparison Between CA199 Assays.* Correlations between the different CA19-9 assays in the training set were assessed through Pearson correlation at natural-log-transformed scale. Different CA19-9 assays were also compared with respect to the classification performance, including AUC, sensitivity at 95% specificity, specificity at 95% sensitivity, and sensitivity and specificity at the 37U/ml threshold. A Wald test was used to determine whether there was a performance difference between assays based on nonparametric bootstrap standard error estimates of assay difference.

### Supplementary Tables

**Supplementary Table 1. Individual Biomarker Performance in the Training Set.** UPMC, University of Pittsburgh Medical Center; VAI, Van Andel Institute; MDACC, MD Anderson Cancer Center; UNMC, University of Nebraska Medical Center; AUC, area-under-the-curve in receiver-operator characteristic analysis; CI, confidence interval in bootstrap analysis.

| Site | Biomarker | AUC (95% CI) | Sens at 95% Spec (95% CI) | Spec at 95% Sens (95% CI) |
|---|---|---|---|---|
| UPMC | TGM2 | 0.27 (0.19, 0.36) | 0.02 (0.00, 0.06) | 0.00 (0.00, 0.03) |
| | THSP2 | 0.68 (0.59, 0.78) | 0.17 (0.05, 0.44) | 0.06 (0.00, 0.29) |
| | HEP | 0.67 (0.58, 0.77) | 0.24 (0.08, 0.39) | 0.11 (0.00, 0.24) |
| | TIMP2 | 0.60 (0.51, 0.69) | 0.21 (0.08, 0.32) | 0.11 (0.00, 0.27) |
| | ANG | 0.44 (0.35, 0.54) | 0.12 (0.03, 0.23) | 0.03 (0.00, 0.09) |
| VAI | CA199.STRA | 0.86 (0.79, 0.93) | 0.61 (0.29, 0.82) | 0.33 (0.02, 0.65) |
| | MUC16.STRA | 0.51 (0.41, 0.61) | 0.18 (0.05, 0.29) | 0.05 (0.00, 0.17) |
| MDACC | TIMP1 | 0.64 (0.55, 0.74) | 0.14 (0.03, 0.39) | 0.03 (0.00, 0.18) |
| | LRG1 | 0.62 (0.52, 0.72) | 0.09 (0.03, 0.27) | 0.08 (0.00, 0.18) |
| UNMC | MUC4 | 0.51 (0.42, 0.62) | 0.09 (0.00, 0.20) | 0.00 (0.00, 0.00) |
| | MUC5AC | 0.53 (0.43, 0.63) | 0.02 (0.00, 0.08) | 0.00 (0.00, 0.12) |
| Clinical | CA199 | 0.82 (0.74, 0.90) | 0.50 (0.27, 0.74) | 0.02 (0.00, 0.61) |
| UPMC | CA199 | 0.86 (0.79, 0.92) | 0.53 (0.35, 0.70) | 0.42 (0.14, 0.71) |
| VAI | CA199 | 0.82 (0.74, 0.90) | 0.47 (0.15, 0.77) | 0.11 (0.03, 0.47) |

| | | | Sens, cutoff 37 U/mL (95% CI) | Spec, cutoff 37 U/mL (95% CI) |
|---|---|---|---|---|
| Clinical | CA199 | - | 0.76 (0.64, 0.85) | 0.79 (0.67, 0.88) |
| UPMC | CA199 | - | 0.92 (0.83, 0.97) | 0.45 (0.33, 0.58) |
| VAI | CA199 | - | 0.68 (0.56, 0.79) | 0.91 (0.81, 0.97) |

**Supplementary Table 2. Classification Rules of Biomarker Panels with corresponding cutoffs determined from the training data.** Logi-Reg, logistic regression; RF, random forest.

| | Basis Biomarker | Additional Biomarkers | Method | Positive Test Definition |
|---|---|---|---|---|
| High specificity | Clinical CA19-9 | - | - | CA19-9 ≥ 164 |
| | Clinical CA19-9 | CA199.STRA | OR | CA19-9 ≥ 375 or CA19-9:STRA ≥ 7587 |
| | Clinical CA19-9 | CA199.STRA + TIMP1 | OR | CA19-9 ≥ 375 or CA19-9:STRA ≥ 7587 or TIMP1 ≥ 399 |
| | Clinical CA19-9 | CA199.STRA + ANG | OR | CA19-9 ≥ 375 or CA19-9:STRA ≥ 7587 or ANG ≥ 288119 |
| | Clinical CA19-9 | CA199.STRA + MUC16.STRA | OR | CA19-9 ≥ 375 or CA19-9:STRA ≥ 7587 or MUC16:STRA ≥ 1171 |
| | Clinical CA19-9 | CA199.STRA + TGM2 + HEP | RF | Predicted case probability ≥ 0.745 |
| | CA199.STRA | ANG | OR | CA19-9:STRA ≥ 7587 or ANG ≥ 288119 |
| | CA199.STRA | MUC16.STRA | OR | CA19-9:STRA ≥ 7587 or MUC16:STRA ≥ 1171 |
| | CA199.STRA | MUC5AC | Logi-Reg | -7.22 + 0.93 * ln(CA19-9:STRA) - 0.09 * ln(MUC5AC + 0.001) ≥ 1.09 |
| | CA199.STRA | THSP2 + MUC5AC | RF | Predicted case probability ≥ 0.828 |
| | CA199.STRA | THSP2 + MUC5AC + ANG | Logi-Reg | -4.27 + 0.88 * ln(CA19-9:STRA) + 0.19 * ln(THSP2) - 0.09 * ln(MUC5AC + 0.001) - 0.40 * ln(ANG) ≥ 1.10 |
| High sensitivity | Clinical CA19-9 | | - | CA19-9 ≥ 2.0 |
| | Clinical CA19-9 | TIMP1 | OR | CA19-9 ≥ 14.0 or TIMP1 ≥ 185 |
| | Clinical CA19-9 | LRG1 | OR | CA19-9 ≥ 13.8 or LRG1 ≥ 124695 |
| | Clinical CA19-9 | THSP2 | OR | CA19-9 ≥ 13.8 or THSP2 ≥ 26417 |
| | Clinical CA19-9 | CA199.STRA + TIMP1 | OR | CA19-9 ≥ 113 or CA19-9:STRA ≥ 1067 or TIMP1 ≥ 185 |
| | Clinical CA19-9 | CA199.STRA + LRG1 | OR | CA19-9 ≥ 113 or CA19-9:STRA ≥ 923 or LRG1 ≥ 124695 |
| | Clinical CA19-9 | CA199.STRA + LRG1 + MUC5AC | Logi-Reg | -16.05 - 0.06 * ln(CA19-9) + 0.97 * ln(CA19-9:STRA) + 0.77 * ln(LRG1) - 0.12 * ln(MUC5AC) ≥ -1.64 |
| | Clinical CA19-9 | CA199.STRA + THSP2 + TIMP1 | RF | Predicted case probability ≥ 0.096 |
| | CA199.STRA | MUC16.STRA | OR | CA19-9:STRA ≥ 923 or MUC16:STRA ≥ 69 |
| | CA199.STRA | LRG1 | OR | CA19-9:STRA ≥ 1444 or LRG1 ≥ 88779 |
| | CA199.STRA | TIMP1 | OR | CA19-9:STRA ≥ 1067 or TIMP1 ≥ 145 |
| | CA199.STRA | THSP2 + MUC5AC | RF | Predicted case probability ≥ 0.106 |
| | CA199.STRA | THSP2 + MUC5AC + ANG | Logi-Reg | -4.27 + 0.88 * ln(CA19-9:STRA) + 0.19 * ln(THSP2) - 0.09 * ln(MUC5AC + 0.001) - 0.40 * ln(ANG) ≥ -1.46 |
| | CA199.STRA | THSP2 + MUC5AC + TIMP1 | RF | Predicted case probability ≥ 0.141 |

**Supplementary Table 3. Individual Biomarker Performance in the Validation Set.** AUC, Area-Under-the-curve in receiver-operator characteristic analysis; CI, confidence interval in bootstrap analysis.

| Lab | Biomarker | AUC (95% CI) | Sens at 95% Spec (95% CI) | Spec at 95% Sens (95% CI) |
|---|---|---|---|---|
| UPMC | TGM2 | 0.73 (0.67, 0.79) | 0.25 (0.07, 0.36) | 0.16 (0.10, 0.31) |
| | THSP2 | 0.67 (0.60, 0.73) | 0.20 (0.11, 0.33) | 0.07 (0.02, 0.23) |
| | HEP | 0.69 (0.63, 0.75) | 0.27 (0.18, 0.36) | 0.11 (0.04, 0.28) |
| | TIMP2 | 0.62 (0.55, 0.69) | 0.16 (0.08, 0.24) | 0.10 (0.04, 0.22) |
| | ANG | 0.53 (0.48, 0.60) | 0.08 (0.02, 0.15) | 0.06 (0.02, 0.14) |
| VAI | CA199.STRA | 0.87 (0.82, 0.90) | 0.59 (0.35, 0.76) | 0.00 (0.00, 0.00) |
| | MUC16.STRA | 0.68 (0.61, 0.73) | 0.29 (0.18, 0.38) | 0.00 (0.00, 0.00) |
| MDACC | TIMP1 | 0.63 (0.57, 0.70) | 0.17 (0.06, 0.29) | 0.05 (0.01, 0.11) |
| | LRG1 | 0.66 (0.59, 0.72) | 0.08 (0.03, 0.19) | 0.18 (0.05, 0.24) |
| UNMC | MUC4 | 0.56 (0.51, 0.62) | 0.12 (0.06, 0.18) | 0.00 (0.00, 0.00) |
| | MUC5AC | 0.71 (0.66, 0.77) | 0.25 (0.14, 0.34) | 0.23 (0.08, 0.37) |
| Clinical | CA199 | 0.84 (0.79, 0.88) | 0.60 (0.47, 0.72) | 0.10 (0.00, 0.30) |
| UPMC | CA199 | 0.86 (0.81, 0.89) | 0.59 (0.51, 0.69) | 0.37 (0.11, 0.54) |
| VAI | CA199 | 0.84 (0.79, 0.88) | 0.59 (0.46, 0.69) | 0.23 (0.11, 0.45) |

| | | | Sens, cutoff 37 U/mL (95% CI) | Spec, cutoff 37 U/mL (95% CI) |
|---|---|---|---|---|
| Clinical | CA199 | - | 0.68 (0.61, 0.75) | 0.90 (0.83, 0.94) |
| UPMC | CA199 | - | 0.96 (0.92, 0.98) | 0.37 (0.28, 0.46) |
| VAI | CA199 | - | 0.65 (0.57, 0.72) | 0.91 (0.85, 0.96) |

### Supplementary Figures

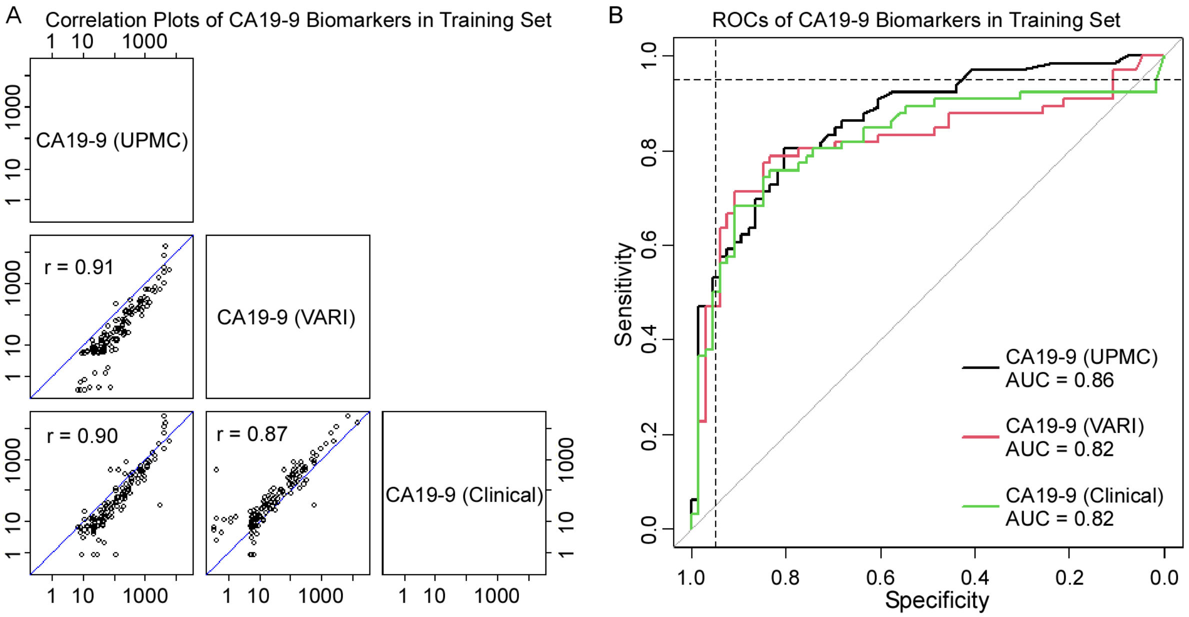

**Supplementary Figure 1. Comparison of Clinical CA19-9 with the Two Research CA19-9 Results.** A) Pearson correlations at natural-log scale. B) Receiver-operator characteristic curves for distinguishing PDAC from all controls.

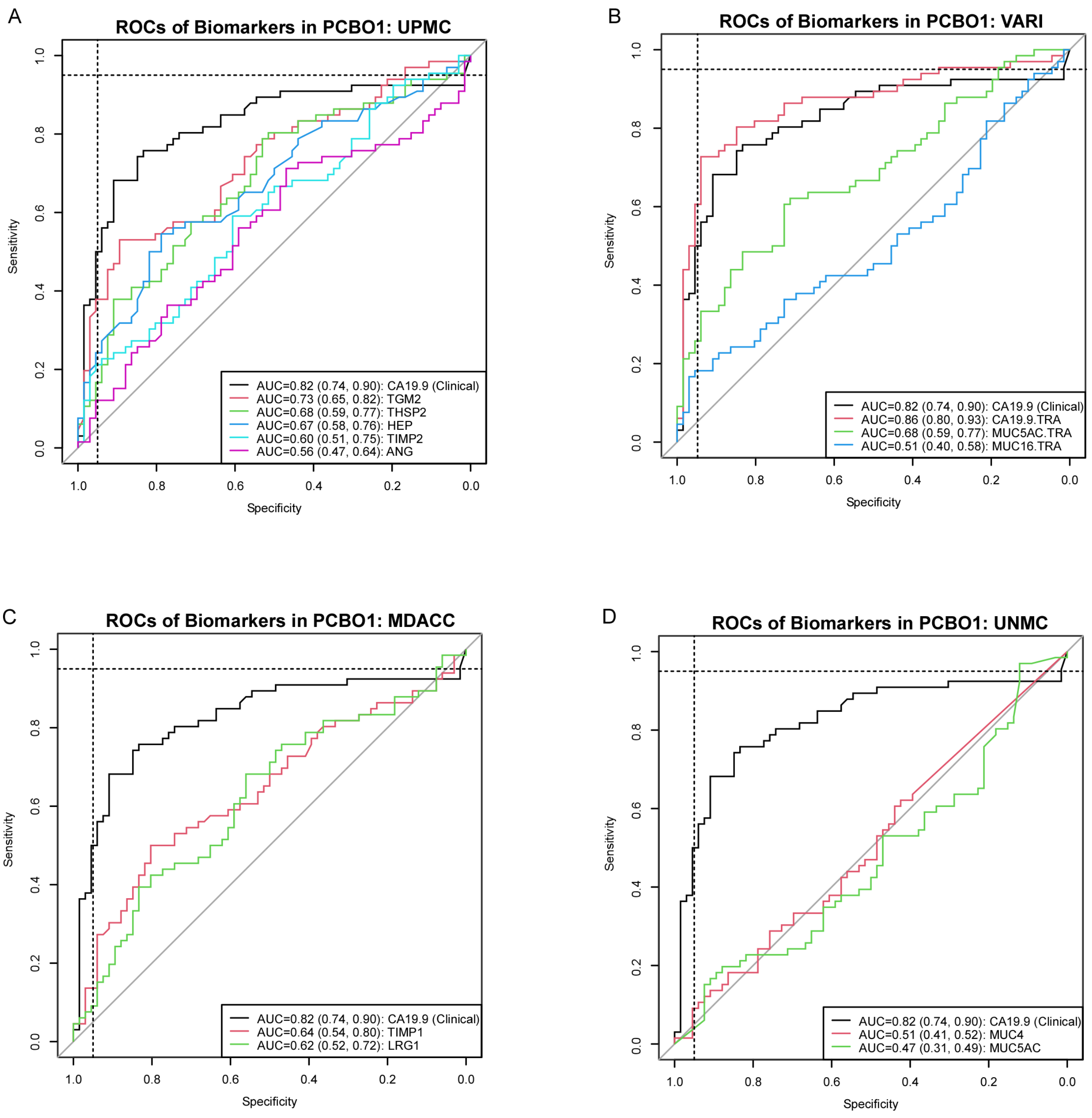

**Supplementary Figure 2. Receiver-Operator Characteristic (ROC) Curves from The Training Set for Each Lab's Individual Biomarkers Compared with Clinical CA19-9.** The ROC curves were generated for distinguishing PDAC from all controls. AUC, area under the curve.

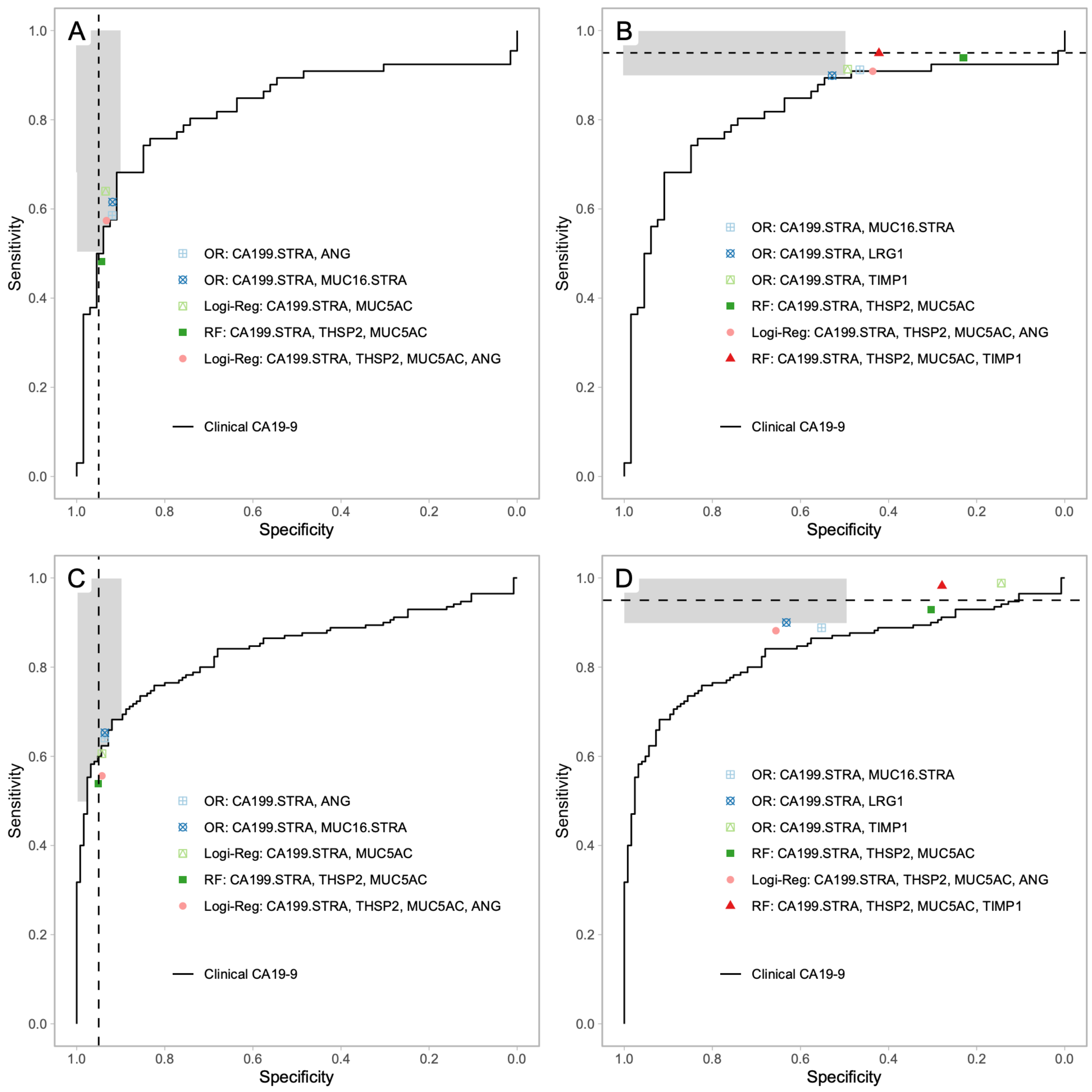

**Supplementary Figure 3. Training Set and Validation Set Performance of Combinations without CA19-9.** The receiver-operator characteristic curves for CA19-9 are shown with the point values of the biomarkers combinations developed for A) high specificity and B) high sensitivity in the training set (with cross-validation) and for C) high specificity and D) high sensitivity in the validation set. The gray boxes indicated the target regions of performance enhancement.
